# Loss of piR-hsa-7221 regulation drives the expression of the LINE1-derived oncogenic lncRNA CASC9 in testicular cancer

**DOI:** 10.64898/2026.02.14.705912

**Authors:** Ahmad Zyoud, Ryan P Cardenas, Nabeelah Almalki, Tinyiko Modikoane, Mohammed Ageeli Hakami, Mansour Alsaleem, Cristina Tufarelli, Nigel P Mongan, Cinzia Allegrucci

## Abstract

Testicular germ cell tumours (TGCTs) are the most common cancer in young males and are considered curable if they respond to platinum-based therapy. However, a significant number of refractory patients develop metastatic disease and the lack targeted therapy remains an unmet clinical need.

To identify novel therapeutic targets, we investigated the epigenetic instability of TGCTs and characterised novel oncogenic gene networks regulated by transposable elements (TEs)-derived long noncoding RNAs (lncRNAs) which are controlled by PIWI-interacting RNAs (piRNAs). A TGCT-specific piRNA signature was identified by bioinformatics analysis of the The Cancer Genome Atlas (TCGA) TGCT dataset and analysis of piRNAs mapped to active LINE1 sequences identified piR-hsa-7221 as a transcriptional regulator of the lncRNA CASC9 in seminoma tumours. We show that piR-hsa-7221 binds to a complementary LINE1 LIPA5 sequence and regulates the expression of CASC9 driven by the LINE1 antisense promoter. Therefore, loss of piR-hsa-7221 drives the upregulation and oncogenic activity of CASC9, which as is impaired after silencing, leading to reduced cancer cell proliferation and invasion, as well as increased sensitivity to cisplatin treatment. These effects are associated with the regulation of the cell cycle, developmental pathways, extracellular matrix, hormone metabolism and immune responses, highlighting WNT signalling as a significant downstream target. Therefore, this novel epigenetic mechanism provides new insights into the role of piRNA-mediated regulation of oncogenic lncRNAs derived from active transposable elements. Importantly, the identification of piR-hsa-7221 and the lncRNA CASC9, together with the associated gene networks highlights novel therapeutic targets for the treatment of seminoma TGCTs.

## Introduction

Testicular germ cell tumours (TGCTs) are the most common type of cancer affecting young men worldwide, with seminoma (SEM) representing the main tumour subtype (1). Despite the increasing incidence reported over the last two decades (2), TGCTs are considered a curable disease if patients respond to conventional cisplatin treatment (3). However, drug resistance can occur in approximately 10–15% of patients with advanced disease and in 3–5% of refractory cases (4, 5). In addition, chemotherapy treatment is highly toxic to germ cells and other organs, compromising the quality of life of cancer survivors (6). Therefore, new research to develop targeted treatments to reduce cancer recurrence and drug toxicity is needed.

TGCTs present significant genomic instability, which contributes to cancer development and progression (7). Epigenetic silencing of transposons is critical to the maintenance of genomic integrity and this process is compromised during cancer evolution, resulting in the aberrant activation of transposable elements (TEs) and consequent genomic rearrangements through retrotransposition and genome integration (8, 9). Among TEs, LINE1 sequences represent the primary source of retrotransposition in the human genome, as they have retained the ability of active mobilisation (10, 11). In addition to impacting on the expression of LINE1 proteins required for retrotransposition and mobilisation of nonautonomous TE, the activity of LINE1 has an important effect on the expression of neighbouring genes through the activity of its bidirectional promoter which can drive the antisense transcripts with oncogenic activity (12, 13).

SEM tumours are characterised by a hypomethylated genome, and a correlation between decreased levels of LINE1 methylation and the development of TGCTs has been reported in patients with SEM (14), supporting the idea that TEs activation could be a driver of testicular cancer. In the germline, LINE1 silencing is mediated by the activity of piRNAs, a class of small noncoding RNAs interacting with PIWI proteins, which can silence TEs through transcriptional and post-transcriptional regulation (15, 16).

The PIWI-piRNA pathway has a critical role in maintaining genomic integrity required for normal spermatogenesis and fertility (17, 18). piRNAs and PIWI proteins are expressed in the male germline throughout the development of primordial germ cells (PGCs), gonocytes and spermatocytes. During the prenatal developmental stages, piRNAs and the associated PIWI proteins PIWIL2 and PIWIL4, recruit DNA methyltransferase and histone methylases to TE sequences (19). In the postnatal stages, silencing of TEs is mostly dependent on the interaction of piRNAs with PIWIL1 and PIWIL2 proteins, which mediate TEs slicing and degradation (18, 20). In testicular cancer, the expression of piRNAs machinery has been found reduced compared to normal testis, together with a decrease in LINE1 methylation (21–23). However, the functional impact of this observation on the disease pathogenesis has not been elucidated. Given that TE sequences are prevalent in lncRNAs and can act as their functional domains (24), in this study we tested whether piRNA dysregulation can drive testicular cancer by affecting the expression of TE-derived lncRNAs. Through the first comprehensive transcriptome analysis of piRNAs expression in TGCTs, we identified a new oncogenic gene network represented by piR-hsa-7221 and LINE1-derived lncRNA CASC9 which drive SEM tumour growth, invasion and drug resistance.

## Methods

### Bioinformatics analysis of piRNAs and lncRNAs expression

Pair-end RNA sequencing (RNA-seq) data of 150 TGCT patient samples from The Cancer Genome Atlas (TCGA) were retrieved after NIH approval (dgGaP Project Approval #12671, Study Accession phs000178). TGCT samples with both clinical and pathological data (n=115) were classified into seminoma (SEM, n=58) and non-seminoma (NON-SEM, n=57) subtypes with a tissue purity of 95%. TGCT RNA-seq data were analysed together with RNA-seq data from 4 publicly available normal testis samples obtained from SRA bioprojects: PRJEB21088 (accession numbers ERX2054613, ERX2054614, ERX2054615) and PRJNA324812 (accession number GSM2193189).

For analysis, the paired-end FASTQ files for forward and reverse reads were concatenated and adaptors were trimmed based on quality (Phred quality score ≥30) and size (read length ≥21 bp) via the publicly available software Trim Galore rapper for Cutadapt and FastQC. The resulting reads were aligned to the human genome (hg38) using the Spliced Transcripts Alignment to a Reference (STAR) software allowing for one mismatch during the remapping of sequence reads (25). A custom piRNA reference transcriptome was created by using the bed coordinates of the piRNAs that were downloaded from the piRBase database v2.0 (26). The coordinates were converted to the human reference genome (hg38/ GRCh38) using the open source UCSC Batch Coordinate Conversion (liftOver) (https://genome.ucsc.edu/cgi-bin/hgLiftOver) together with the UCSC utilities BedToGenepred and GenepredToGTF. The piRNA file was subsequently concatenated with a GENCODE annotation file (version 25, GRCh38) to produce the final custom annotation. Finally, the piRNA read counts were obtained by annotating the samples aligned to the custom piRNA reference transcriptome using FeatureCounts, with piRNA genomic coordinates obtained from the piRBase database (27). Counts of identical piRNA sequences with different genomic loci were merged using the Kutools add-in for Microsoft Excel to aggregate all the identical sequence counts into one ID using R (28). Reads were filtered by applying a threshold of count >1 in at least 10% of the samples by using the open source software Cluster 3.0 (29) and then analysed with the R based software RobiNA with the edgeR method and the Benjamini and Hochberg correction of False Discovery Rate (FDR) (30) to obtain differentially expressed piRNAs (FDR<0.05). Differentially expressed piRNAs were overlapped with TE element transcripts retrieved from the USCS repeat masker feature and lncRNA transcripts obtained from the GENCODE database using Galaxy (31).

Expression heatmaps, correlation matrices and dendrograms were created via the R software (version 4.0) after data counts were converted to transcript per million (RPKM) and filtered using the Cluster 3.0 software (29) to remove transcripts with a difference between maximum and minimum read values < 100. Hierarchical clustering of the heatmap was performed using the Euclidean distance of transcript expression and the Ward’s method. Dendrograms were obtained using the Bootstrap method from 1000 replicates (32), with clusters considered significant when the approximately unbiased (AU) probability value was > 95%.

Circular representation of genome-wide piRNAs expression in TGCTs was performed using CircosPlot (33) after log transformation of the read values and filtration to remove transcripts with values between 1 and -1 RPKM using Cluster 3.0 (29). Volcano plots, box plots and histograms were created with the GraphPad Prism software. Venn diagrams were created with Venny 2.0.2 (https://bioinfogp.cnb.csic.es/tools/venny/index.html).

### RNAseq analysis

Total RNA for RNAseq analysis was extracted using a miRNeasy kit (Qiagen). A total of four biological replicates per sample were processed and analysed by Novogene Co., Ltd (Cambridge, UK). Before sequencing, the RNA Nano 6000 Assay Kit and the Bioanalyzer 2100 system (Agilent Technologies, CA, USA) were used to assess RNA integrity. The sequencing libraries were generated using the NEBNext® UltraTM RNALibrary Prep Kit for Illumina® (NEB) using index codes that were clustered via a cBot Cluster using a PE Cluster Kit cBot-HS (Illumina). The samples were then sequenced using the Illumina 6000 S4 platform with pair-end reads (150 bp). Clean reads without adapters were obtained via fastp, with Q20, Q30, and GC contents calculated to ensure high quality data (QC 93.39-98.06%; error rate= 0.03%). Paired-end clean reads were aligned to the reference genome (GRCh38/hg38) using the Spliced Transcripts Alignment to a Reference (STAR v2.5) software (mapping rate 95.9-97.5%). Reads were quantified with FeatureCounts (34) and normalised to gene length (FPKM counts). The RNAseq data are available from the GEO database under accession number GSE232791. Differential expression was performed via DESeq2 (v2_1.6.3) with the FDR set at < 0.05 for significant differences. Gene Ontology (GO), GSEA and pathway analyses were performed using the software Webgestalt (35).

### Cell culture and transfections

The human testicular seminoma germ cell tumour TCam-2 cell line (CVCL-T012) (36) was kindly donated by Prof. Azim Surani (University of Cambridge). Cells were cultured in RPMI 1640 medium supplemented with 10% foetal calf serum, 1% glutamine,1% MEM nonessential amino acids, and 1% sodium pyruvate (reagents from ThermoFisher) and incubated at 37°C with 5% of CO_2_. TCam-2 cells were routinely tested for mycoplasma using the PlasmoTest™ (InvivoGen) and genotyped using the Applied BiosystemsTM AmpFLSTRTM IdentifilerTM Plus PCR Amplification Kit system (Eurofins Genomics).

For luciferase analysis, TCam-2 cells were seeded in a 24-well plate at a density of 50,000 cells per well and the psiCHECK-2 plasmid (Promega C8011) containing the LINE1 promoter sequence was co-transfected with piRNA mimics (scrambled or piR-hsa-7221, 25 nM) and plasmids encoding for full-length PIWI proteins (PIWIL1, PIWIL2, and PIWIL4; plasmids modified from Addgene plasmid # 16578) using the TransIT-X2® transfection reagent (Mirus) (Table S1). Cells were analysed 3 days after transfection.

To study the function of piR-hsa-7221, TCam-2 cells were seeded in a 12-well plate at a density of 300,000 cells per well. The PIWI plasmids (PIWIL1, PIWIL2 and PIWIL4 were co-transfected with piRNA mimics (scrambled or piR-hsa-7221, 25 nM) twice (3 days apart) and cells were collected on day 6.

For CASC9 silencing, a mixture (1:1) of the Silencer® Select CASC9 siRNAs (assays # n544070 and n544071; ThermoFisher) was transfected at a final concentration of 25 nM with the TransIT-X2® transfection reagent (Mirus). Cells were analysed on day 4 after transfection.

### Cell proliferation, invasion and drug response

The MTT (3-(4,5-dimethylthiazol-2-yl)-2,5-diphenyltetrazolium bromide) assay was used to measure cell viability. Cells were incubated with 5 µg/ml MTT for 3 hours at 37°C and then treated with acidic isopropanol (0.04 M hydrochloric acid in absolute isopropanol) to solubilise the converted formazan dye. The spectrophotometric absorbance was measured at a wavelength of 570 nm with 650 nm background subtraction.

For cisplatin treatment, cells were seeded at 3,000/well in a 96-well plate and transfected with CASC9 or scrambled siRNAs. Sensitisation to cisplatin was assessed 3 days after transfection and after 72 hours of treatment with 25 µM and 50 µM (IC25 and IC50, respectively as previously described (37).

Cell invasion was measured by first transfecting the cells with siRNAs for 3 days and then plating 2.5×10^5^ cells into a Falcon cell culture insert (8.0 µm pores) coated with 300 µg/ml Matrigel (Corning) in serum-free RPMI medium and fitted into a companion plate. RPMI medium supplemented with 10% FCS was placed at the bottom of the plate to allow for cell invasion, which was assessed after 24 hours of incubation and after the removal of the cells from the top insert compartment. Cells were fixed with 100% methanol and stained with 0.4% crystal violet in 10% methanol. A Nikon Eclipse TE2000-S inverted microscope was used to image the cells with the Nikon NIS-Elements software.

### RT-qPCR

The miRNeasy Mini Kit (Qiagen) was used to extract RNA which was then reverse transcribed with a stem-loop method using the TaqMan™ MicroRNA Reverse Transcription Kit (ThermoFisher) for piRNA analysis or using the Superscript IV (ThermoFisher) for mRNA analysis. The stem-loop sequence and annealing conditions were designed as previously described (38). RT-qPCR was performed using the LuminoCt® SYBR® Green qPCR ReadyMix (Sigma Aldrich). The ΔΔCT method was used to analyse the data that were normalised to the endogenous control genes U6 snRNA and ACTB for piRNA and mRNA gene expression, respectively. Endogenous control genes were selected after assessment of expression stability using the software BestKeeper (39). Total RNA from normal testes was obtained from Clontech (# 636533). The primers utilised are listed in Table S1. CASC9 primers were designed to amplify different transcript variants, as indicated.

### Bisulfite sequencing

For methylation analysis, genomic DNA was extracted using the DNeasy Blood & Tissue Kit (Qiagen), and 1µg of DNA was then converted by bisulfite reaction for 16 hours at 50°C using the EZ DNA Methylation Kit (Zymo Research). Converted DNA was purified using a QIAquick PCR & Gel Cleanup Kit (Qiagen) and treated with 3M sodium hydroxide at room temperature for 20 minutes, followed by neutralisation with sodium acetate. The converted DNA was further purified using the same kit. For Bisulfite sequencing cloning, the region of interest was amplified using the Platinum Taq DNA polymerase (ThermoFisher), gel purified and then cloned into the pGEM®-T Easy Vector (Promega) with white/blue screening. Plasmid DNA was purified with the QIAprep Spin Miniprep Kit (Qiagen) and at least 10 clones were sequenced by Sanger sequencing (Source BioScience). The methylation primers are listed in Table S1.

### Western blotting and immunocytochemistry

Whole-cell lysates were prepared using the cOmplete™ Lysis-M EDTA-free reagent (Merck). Protein concentrations were quantified using the Pierce BCA kit assay (ThermoFisher) and 30 μg of the protein was separated on SDS-bolt bis-tris gels (4–12%) before being transferred onto a nitrocellulose membrane. The membrane was blocked with 5% skimmed milk and then probed overnight at 4°C with mouse anti-L1-ORF1p monoclonal antibody (Merck Millipore #MABC1152, 1:1,000) and rabbit anti-ACTB (Cell Signalling #4970, 1:1,000). The secondary antibodies were incubated for 1 hour at room temperature (Li-COR IRDye 800CW Donkey Anti-Rabbit IgG #925-32211 and IRDye 680CW Donkey Anti-Mouse IgG # 926-68072 at a dilution of 1:10,000). The membrane was imaged using an Odyssey XF Imager (Li-COR).

For immunocytochemistry, TCam-2 cells were grown on coverslips and fixed with 4% paraformaldehyde for 15 minutes, permeabilised with 0.1% Triton™ X-100 for 10 minutes and blocked with 3% BSA-PBS for 30 minutes at room temperature.

Cells were then incubated with mouse anti-L1-ORF1p monoclonal antibody (Merck Millipore MABC1152, 1:1, 000) in 0.1% BSA-PBS, overnight at 4°C, and then labelled with goat anti-Mouse IgG (H+L) Cross-Adsorbed Secondary Antibody, Cyanine3 (ThermoFisher Scientific, A10521) and Phalloidin Alexa Flur 488 (ThermoFisher # A22282, 1:1,000) for 45 minutes at room temperature. The samples were mounted with Fluoroshield with DAPI and imaged using a Leica TCS SPE confocal microscope (scale bar = 25 μm).

## Statistical analysis

Statistical analyses were performed using the GraphPad Prism software (statistical significance set at P<0.05). Data are presented as the mean ± standard deviation (SD) of biological replicates (≥3 biological replicates in technical triplicates). One-way and two-way ANOVA tests were used to compare more than two groups, whereas two-tailed unpaired Student’s t-test or Mann Witney test were used to compare two groups. Different categories comparisons were analysed by Chi-square test.

## Results

### Analysis of piRNAs expression in TGCTs

To analyse the expression of piRNAs in TGCTs, the RNA-seq raw counts from the TCGA TGCTs and publicly available normal testis (40, 41) datasets were mapped using a custom piRNA transcriptome annotation. A total of 812,382 annotated sequences were analysed, 93,237 of which (11.48%) were identified in normal testis. To assess the accuracy of our custom annotation, sequences were matched to 32,047 piRNAs already identified in available human testis datasets from the piRBase and piRNAdb repositories (26, 42). The sequences expressed in normal testes corresponded to 8,039 unique piRNAs, and only a small minority (0.1%) were identified with a different identity, thus validating the piRNA annotation (Fig. S1). In TGCTs, the majority of these piRNAs were not expressed, and only 1,891 piRNAs were detected (Fig. 1A).

**Fig. 1.**
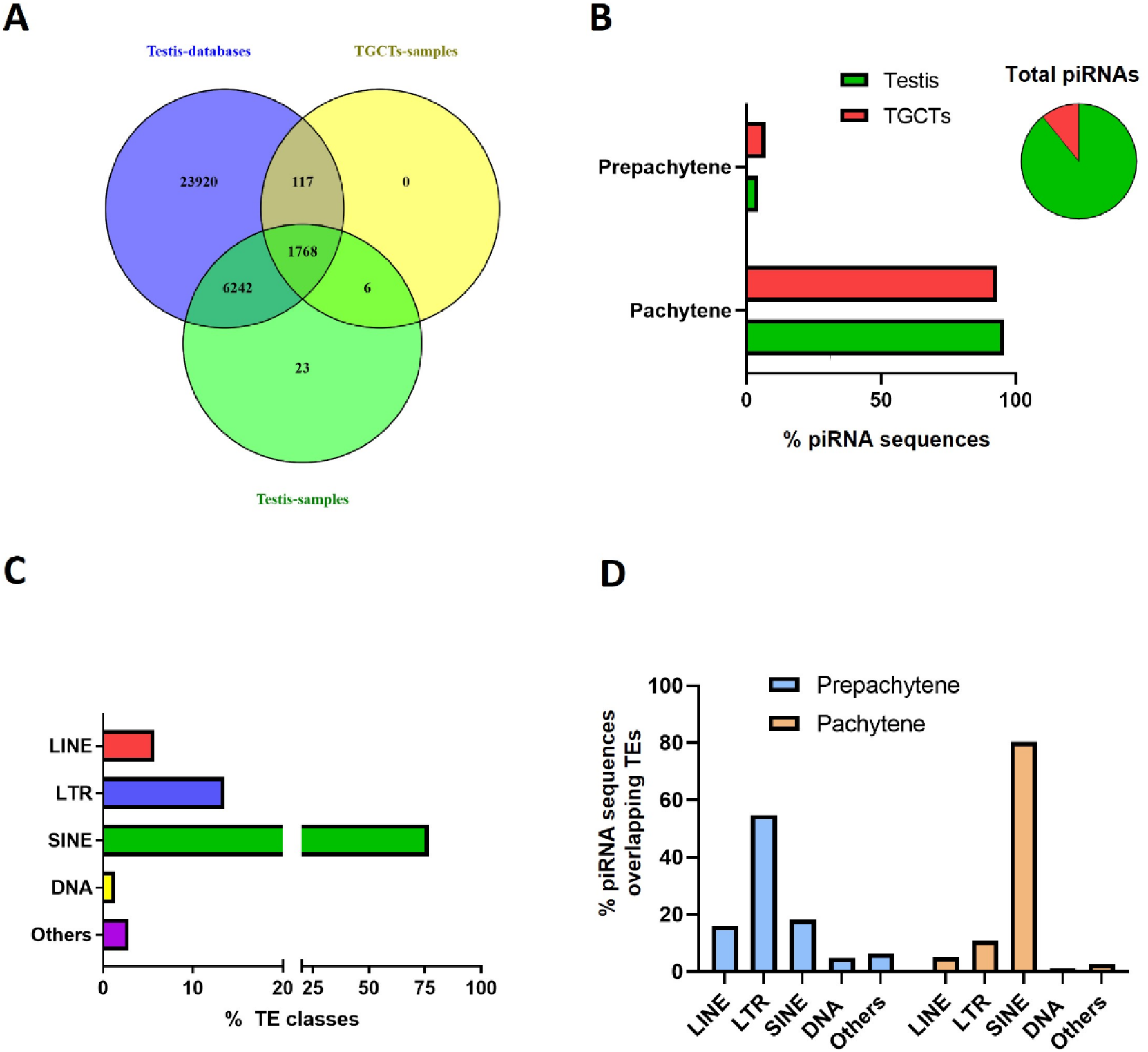
TGCTs piRNA expression signature and overlap with TEs. **A.** Overlap of piRNA sequences from piRBase and piRNAdb identified in TGCTs and normal testis. **B.** Number of TGCT piRNA sequences identified as prepachytene and pachytene. **C.** TE sequence classes overlapping with TGCT piRNA sequences. **D.** Distribution of TE sequences overlapping prepachytene and pachytene piRNA sequences.

We next classified the piRNAs into prepachytene and pachytene according to their sequence (prepachytene less than 26bp; pachytene greater than 27bp) (16). The majority of piRNAs in normal testes and TGCTs were found to be pachytene, which represented 95.7% and 93% of the total number expressed, respectively (Fig 1B). Among the TGCT piRNAs, 4,127 (36.5%) were found to overlap with different types of TE targets (LINE, SINE, LTR, DNA transposons and others) in an antisense orientation, with multiple TEs overlapping the same loci. The majority of TGCT piRNAs overlapped with SINEs, followed by LTRs and LINEs. DNA transposons and other TE classes constituted the minority (Fig. 1C). In total, 252 prepachytene and 3,875 pachytene piRNA overlapped with TEs, with LTRs being predominant in prepachytene and SINEs predominant in pachytene sequences (Fig. 1D).

After the initial sequence mapping, data were further processed to provide a final list of piRNA transcripts by excluding reported piRNAs of ambiguous identity (43) and by binning identical piRNA transcript counts mapped to different genomic loci.

Unsupervised hierarchical clustering of piRNA transcripts was next performed to compare the transcriptional signatures of normal testis and TGCTs classified according to tumour subtype (SEM and NON-SEM). The analysis revealed a distinct clustering of normal testis samples from TGCTs, but no significant clustering was observed between TGCTs clinical subtypes. Likewise, a Spearman correlation-based matrix using rank-normalised piRNA expression levels in TGCTs and normal testes confirmed a high level of correlation for piRNA expression in normal tissue and only a moderate degree of correlation for SEM and NON-SEM tumours (Fig. S2).

The genomic distribution of piRNAs expression after normalisation to normal testis mapped to all chromosomes, with the majority displaying a reduced expression in tumours samples. Interestingly, a significantly lower piRNAs distribution was detected on the X chromosome than on autosomes after normalisation to chromosome length (Chi-square test, p-value <0.05), but no difference in distribution was detected on the Y chromosome. The overall distribution of piRNA expression across all chromosomes was similar between the TGCT subtypes (Chi-square test, p-value >0.05) (Fig. 2A). Therefore, a differential expression analysis was next undertaken to identify piRNAs uniquely expressed in TGCTs, which revealed 471 piRNAs expressed at low level and 172 expressed at high level in tumour samples compared to normal testis (Fig. 2B). These represented a high number of piRNAs compared to other cancer types, which were also expressed at low level, a characteristic shared only with head and neck cancer (Fig. 2C, D). Interestingly, only few piRNAs were shared between TCGTs and other cancers (Table S2), highlighting the tissue-specific expression and function of this class of noncoding RNAs.

**Fig. 2.**
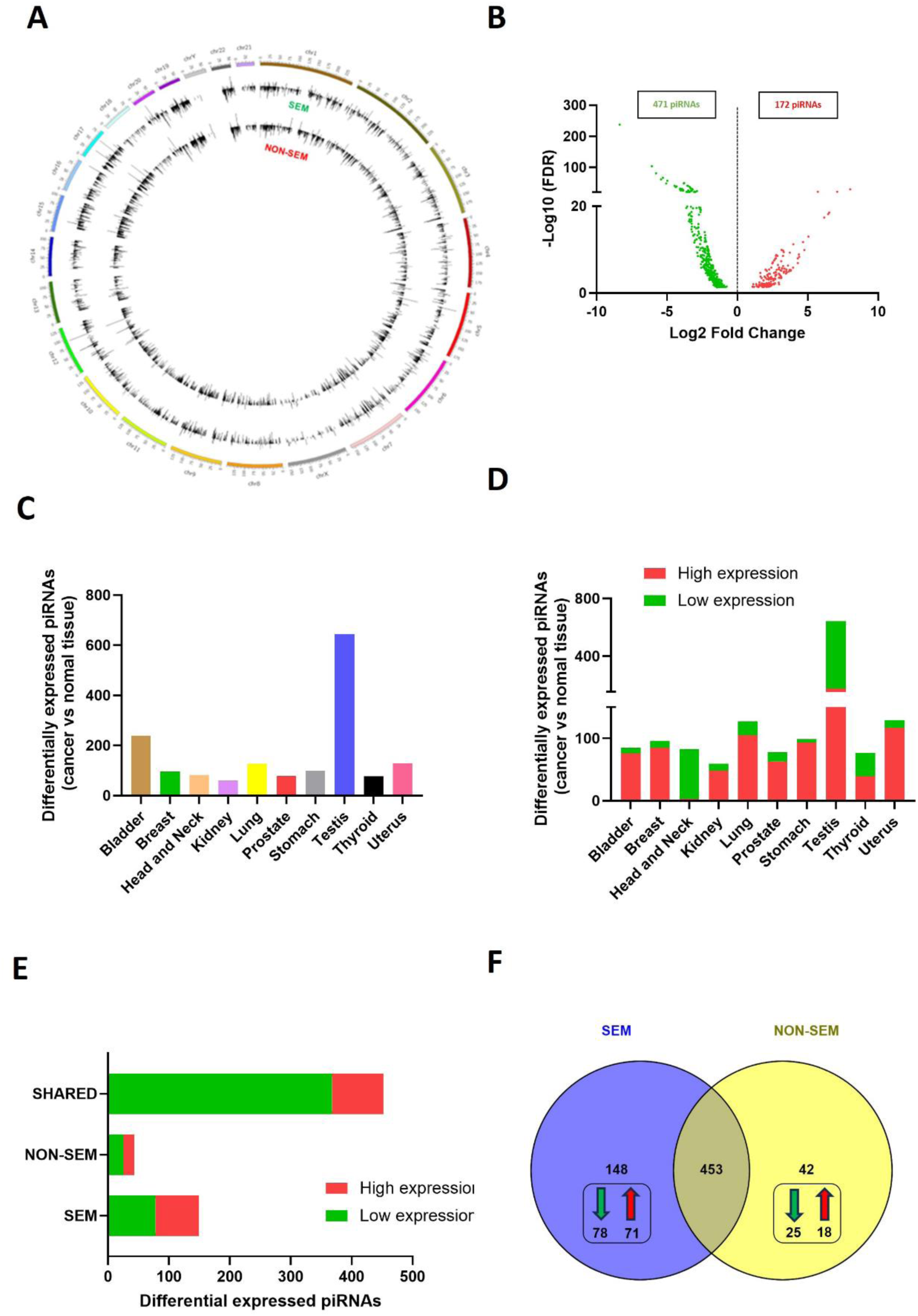
Expression of piRNAs in TGCTs and other cancer types **A.** Circos plot of genome-wide expression of piRNAs in TGCTs. Concentric tracks in the ideogram represent histograms of logarithmic average transcripts of SEM (n=58) and NON-SEM (n=57) normalised to normal tissue expression. **B.** Volcano plot of differentially expressed piRNAs in TGCTs compared to normal testis. **C.** Expression of piRNAs in TGCTs and other cancer types compared to the normal tissue counterparts. **D.** Number of piRNAs expressed at low and high levels in different cancer types. **E.** Differentially expressed piRNAs expressed at high and low levels in SEM (n=58) and NON-SEM (n=57). **F.** Overlap of differentially expressed piRNAs in SEM (n=58) and NON-SEM (n=57). Green and red arrows indicate the number of unique piRNAs expressed at low and high level, respectively.

Finally, to identify piRNAs specific to SEM tumours, a differential expression analysis showed 84.34% piRNAs shared between tumour subtypes (453 piRNAs), whereas 148 and 42 piRNAs were differentially expressed uniquely in SEM and NON-SEM samples, respectively. The majority of shared and subtype specific piRNAs were expressed at low level (Fig. 2E, F), consistent with the data obtained from the genomic distribution analysis (Fig. 2A).

### piRNAs map to LINE1-derived lncRNAs

To confirm our hypothesis that downregulation of piRNA silencing can lead to active LINE1 sequences and expression of LINE1-derived lncRNAs, we next focussed on those piRNAs uniquely expressed at low level in SEM (Fig.3A). To this end, we examined candidate piRNAs with sequence complementarity to TEs overlapping lncRNA transcripts. The analysis identified 12 candidate piRNAs, which corresponded to 175 lncRNA transcripts (Fig.3A, Fig.S3). These lncRNAs were then overlapped with lncRNAs expressed in SEM tumours using the TGCTs TCGA dataset, resulting in 17 candidate lncRNAs, 10 of which found complementary to LINE1 sequences (Fig. 3B). From these SEM-specific candidate transcripts, the lncRNA CASC9 was selected based on its high expression (median expression of Log2 RPKM ≥2) (Fig. 3B).

**Fig. 3.**
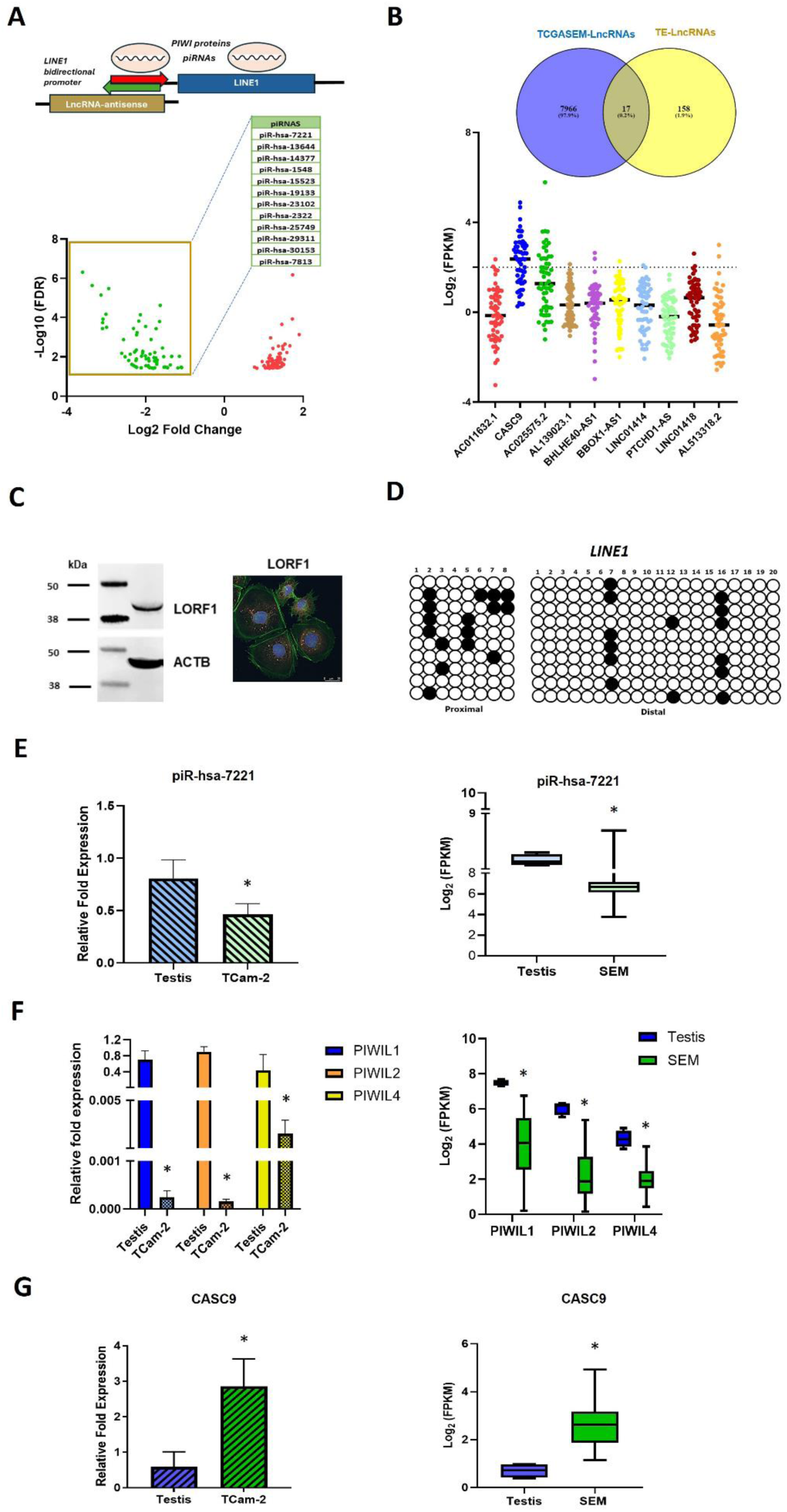
Mapping of piRNAs, LINE1 and lncRNA sequences. **A.** Schematic of piRNAs mapping to lncRNAs transcribed from LINE1 antisense promoters with volcano plot of differentially expressed piRNAs in SEM. IDs of low expressed piRNAs mapping lncRNAs are indicated. **B.** Overlap of LINE1-derived lncRNAs with highly expressed lncRNA in TCGA SEM samples. Expression of overlapping lncRNAs is shown with a cut-off expression of RPKM ≥2 (dotted line). **C.** Western blotting and immunostaining showing expression of LINE1 ORF1 protein (LORF1p) in TCam-2 cells. Expression of ACTB was used as a loading control (LORF1p red, cell cytoskeleton green, nucleus blue). **D.** Bisulfite sequencing of the proximal and distal promoter region of the CASC9-LIPA5 sequence. Ten independent sequence clones are shown with white and black circles representing unmethylated and methylated CG residues, respectively. **E.** PiR-hsa-7221 expression in TCam-2 cells (top; *P < 0.05, Student’s t-test) and in SEM (bottom; TGCA, n= 58; *FDR < 0.05) compared to normal testis. **F.** Expression of PIWIL1, PIWIL2 and PIWIL4 in TCam-2 cells (left; n=3, *P < 0.05, Student’s t-test) and in the TCGA SEM samples (right; n=58, *FDR < 0.05). **G.** CASC9 expression in TCam-2 cells (top; *P < 0.05, Student’s t-test) and in SEM (bottom; TGCA, n= 58; *FDR < 0.05) compared to normal testis.

The visualisation of the CASC9 transcripts at the chr8:75,324,251-75,324,280 locus confirmed the antisense alignment to a LINE1-L1PA5 sequence, which showed complementarity to piR-hsa-7221 (GeneCards ID: piR-45012-495) (Fig.S3).

To validate our bioinformatics screen, we next used the well-characterised SEM cell line TCam-2. To confirm its validity as an *in vitro* model for SEM, we first assessed the methylation and activity of LINE1 promoters in these cells. In agreement with a decrease in piRNAs in SEM, we found that these cells expressed the LINE1 ORF1 protein (LORF1p; Fig. 3C) and that LINE1 promoters are largely unmethylated as bisulfite sequencing of the proximal and distal LINE1 promoter sequences revealed only 20% and 7% methylated CG residues, respectively (Fig. 3D). piR-hsa-7221 and PIWI proteins expression was also confirmed to be lower in TCam-2 cells than normal testis, consistent with the expression in SEM tumours from the TGCTs TCGA patient cohort (Fig. 3E, F). On the other hand, CASC9 expression was higher than normal testis (Fig. 3G), with the CASC9 long transcript variants overlapping the LINE1-L1PA5 sequence and the piR-hsa-7221 binding sequence being the most expressed isoforms (Fig. S3). Therefore, these results confirm an inverse correlation between the expression of piR-hsa-7221 and the LINE1-derived lncRNA CASC9.

### Functional regulation of LINE1-derived CASC9

To study the regulation of the expression of LINE1-derived CASC9, we first tested the binding of piR-hsa-7221 to the LINE1 sequence by transfecting the piRNA mimic and plasmids encoding for PIWI proteins, to restore the expression of the piR-hsa-7221 and PIWIL1, PIWIL2, and PIWIL4 (Fig. 4A, B), together with a psiCHECK-2 plasmid carrying the LINE1 complementary sequence into TCam-2 cells. A decrease in luciferase activity was measured upon the transfection of piR-hsa-7221, which was not affected by the co-transfection of PIWI proteins (Fig.4C), confirming the piRNA binding to LINE1 through base-pairing. To investigate the functional relationship between the expression of piR-hsa-7221 and CASC9, we next expressed a piRNA mimic and PIWI proteins. The expression of piR-hsa-7221 induced a significant reduction of CASC9 expression, which in this case required the concomitant expression of the PIWI proteins PIWIL1, PIWIL2 or PIWIL4. Of note, CASC9 expression downregulation was more pronounced after transfection of PIWIL1 and PIWIL2 compared to PIWIL4 (Fig. 4D).

**Fig. 4.**
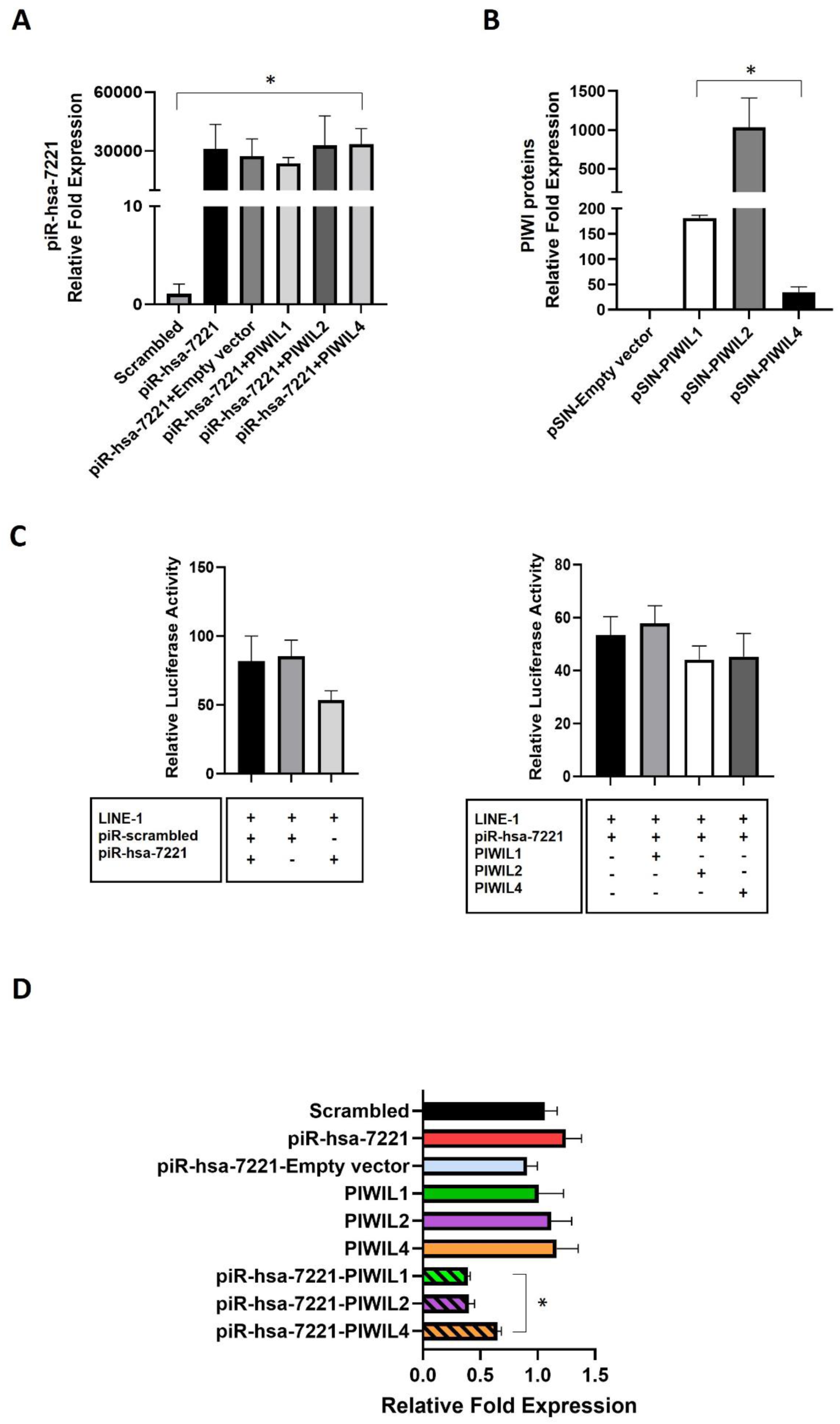
Regulation of LINE1 activity in TGCTs. **A.** Validation of the expression of piR-hsa-7221 after transfection in the presence or absence of PIWI proteins (n = 3) (*P < 0.05, Two-way Anova followed by Tukey’s multiple comparisons test). **B.** Gain of function expression of PIWI proteins in TCam-2 cells. A. Validation of the expression of PIWIL1, PIWIL2, and PIWIL4 after transfection (n = 3) (*P < 0.05, Student’s t-test). **C.** Luciferase assay demonstrating the binding of piR-hsa-7221 to the complementary LINE1 sequence in the presence or absence of co-transfected PIWI proteins. **D.** Regulation of CASC9 expression by piR-hsa-7221 and PIWI proteins (n=3, *P<0.05 compared to piR-hsa-7221, Two-way Anova followed by Tukey’s multiple comparisons test).

### CASC9 oncogenic activity in TGCTs

To evaluate the role of CASC9 in TGCT pathogenesis, we next silenced its expression in TCam-2 cells (Fig. 5A). CASC9 silencing resulted in a significant decrease in cell proliferation measured over 4 days after the siRNA transfection (Fig. 5B). In addition, a significant decrease in cell invasion was observed (Fig. 5C), as well as cell sensitisation to cisplatin, which was increased by ∼12% and 27% at the IC25 and IC50 concentrations of the drug, respectively (Fig. 5D).

**Fig. 5.**
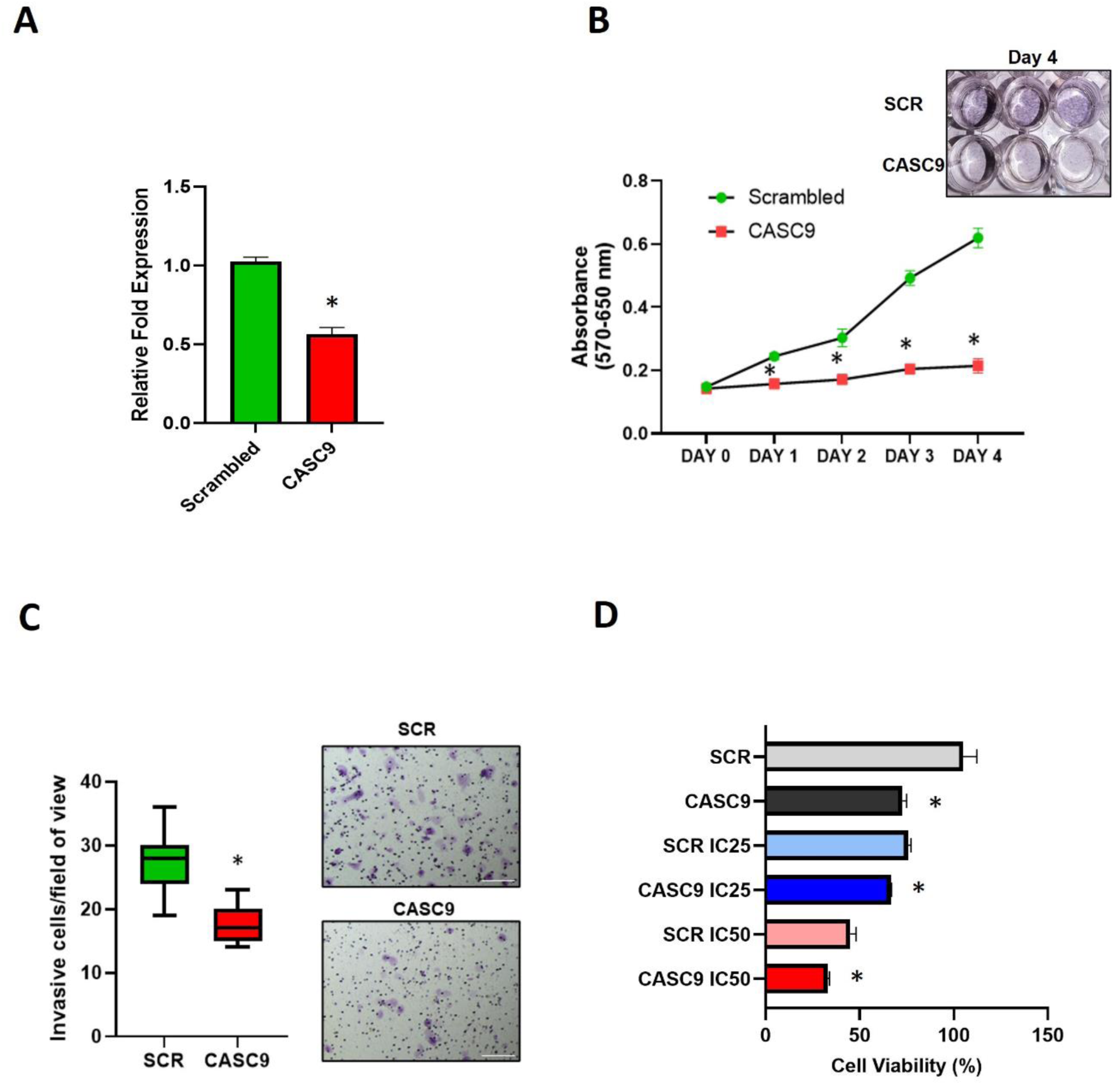
Regulation of CASC9 expression and associated cancer phenotypes. **A.** CASC9 expression after siRNA silencing (n=3, *P < 0.05, Student’s t-test compared to scrambled). **B.** Effect of CASC9 silencing on cell proliferation and cell viability (n = 3; *P < 0.05 compared to scrambled, Two-way Anova followed by Tukey’s multiple comparisons test). Viable cells are stained in purple. **C.** Effect of CASC9 silencing on cell invasion (n = 3, 10 fields of view; *P < 0.05 compared to scrambled, Mann Whitney test), scale = 500 μm; **D.** Effect of CASC9 silencing on cell sensitivity to cisplatin (n = 3; *P < 0.05 compared to scrambled, Two-way Anova followed by Tukey’s multiple comparisons test).

To gain further mechanistic insights into the role of CASC9, we performed RNAseq after CASC9 silencing, which revealed a change in 322 upregulated and 629 downregulated genes (FRD <0.05) (Fig. 6A). Gene ontology analysis revealed associations with extracellular matrix (ECM) structure organisation, cell movement, hormone metabolism, regulation of embryonic development and growth factor binding (Fig. 6B). The most enriched pathways were related to ECM interactions, epithelial to mesenchymal transition (EMT), angiogenesis, myogenesis, hormone metabolism and pathways in cancer (Fig. 6C). Several genes encoding ECM components, including COL6A1, COL18A1, COL5A2, COL6A2, COL1A2, COL22A1, COL3A1, LAMC3, LAMB1, FNB3, DMD, SDC1, FBLN1, HAPLN1, HSPG2, and VIT were downregulated. Similarly, the expression of several genes involved in cell migration, such as CARMIL2, CAPG, NPNT, PHLDB2, HPN, LCP1, PHLDB2, CDH1, MMP15, ADAM15, CXCL12, CX3CL1, CEACAM1 and CLDN4 was reduced. CAPN6 expression was also downregulated with concomitant downregulation of the vascular genes PDGFA, VEGFB, and AMOT (Fig. S4).

**Fig. 6.**
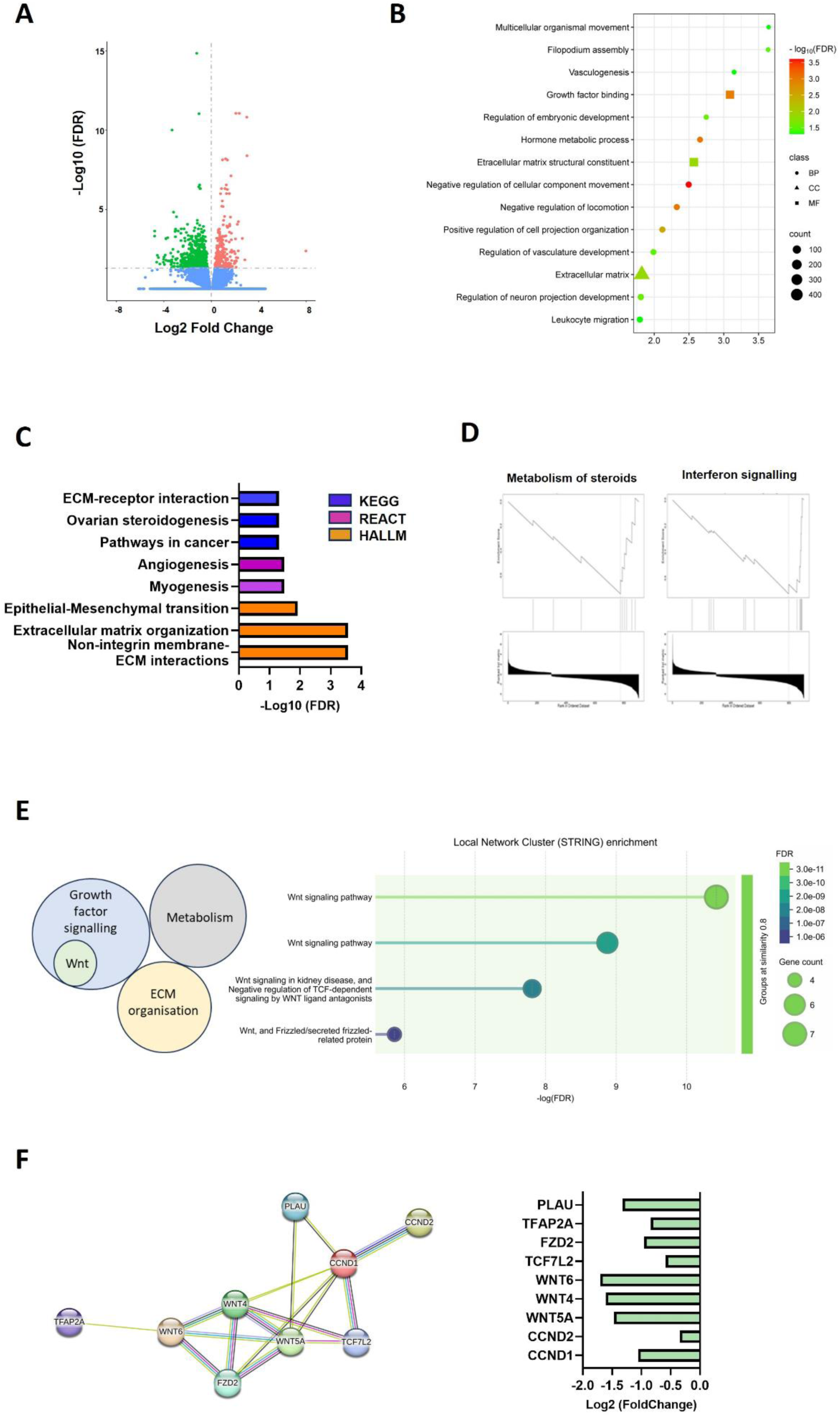
Gene expression signatures enriched in CASC9 knockdown cells (FDR < 0.05, compared to scrambled). **A.** Volcan plot of differentially expressed genes after CASC9 silencing. Green dots represent downregulated genes; red dots represent upregulated genes. **B.** Gene ontology analysis of differentially expressed genes (BP indicates biological process, CC indicates cellular location and MF indicates molecular function). **C.** Enriched pathways. **D.** Enriched GSEA signatures. **E.** Enriched functional protein networks. **F.** WNT signalling protein network cluster and expression fold changes of genes associated with the WNT pathway (FDR<0.05).

Downregulation of transcription factors and enzymes involved in testicular steroidogenesis was also observed, including FOXO4, FOXO6, FOXA3, CYP11A1, CYP19A, HSD3B1, HSD11B1L, CGA, LRP2, and PTGIS. In addition, the metabolism of steroids was among the downregulated GSEA signatures, together with interferon signalling and inflammatory responses (Fig. 6D). The genes included in these signatures were BST2, IFI27, IFI6, MX1, MX2, OAS3, UBE2L6, C3, BTK, PTGES, ADORA1, APOL2, MRGPRX1and TREM1.

Genes involved in cell proliferation, including cell cycle regulators (CCND1, CCND2, CCNG2, CDK18, TFDP2 and TOP2A) were also downregulated. On the other end, cell cycle inhibitors were upregulated (CDK2AP1, GADD45A).

WNT signalling was highly enriched among the downregulated developmental pathways, with GATA2, GATA3, GATA3-AS1, WNT5A, WNT4, WNT6, TCF7L2, FZD2, FZD4, INSR, TFAP2A, TCL1A, SERPINF1, PLAU, PLCB2, and KERMEN2 significantly expressed at low level (Fig. S4). Importantly, WNT signalling was also found as the most significant pathway among the downregulated functional protein association networks (Fig. 6E and Fig.S5). Therefore, taken together these data confirm the role of piRNAs and LINE-1-derived lncRNAs gene regulatory networks as major players in the pathogenesis of TGCTs and establish the oncogenic effect of CASC9 through regulation of WNT signalling, tumour metabolism and microenvironment, thus supporting its role as a possible therapeutic target for the treatment of SEM.

## Discussion

Understanding the molecular mechanisms underlying the pathogenesis of testicular cancer is critical to the development of targeted therapies, which are needed to overcome drug toxicity, resistance, and disease relapse. To explore testicular cancer vulnerabilities, here we investigated a fundamental mechanism involved in the epigenetic regulation of genomic stability in the male germline. The development of germ cells depends on an epigenetic reprogramming that is required to re-establish totipotency and reset parental epigenetic memory via erasure and subsequent re-establishment of DNA methylation and histone methylation marks. The piRNA-PIWI machinery plays a pivotal role in the regulation of this process, which needs to be tightly regulated to evade the activation of TEs for the maintenance of genomic integrity (18).

Through the first comprehensive analysis of TGCTs samples from the TCGA patient cohort, our study shows that the expression of piRNAs and PIWI protein is significantly reduced in testicular cancer compared to normal testis, confirming previous findings from limited studies of testicular tumours and TGCT cell lines (21–23). Silencing of TEs is dependent on the activity of piRNAs during germ cell development. This is mediated either by transcriptional or post-transcriptional regulation, which occurs in partnership with specific PIWI proteins. During the fetal development, piRNAs prepachytene piRNAs facilitate TEs silencing via DNA and histone methylation, whereas both prepachytne and pachytene drive direct TE degradation in postnatal germ cells (16, 18).

Although the majority of piRNAs identified in TGCTs were pachytene, a higher number of prepachytene piRNAs were identified compared to normal testis, suggesting a possible fetal cell of origin for these tumours. When compared to other tumour types, TGCTs presented the highest number of piRNAs, consistent with their prominent role in the germline compared to somatic tissues (25).

Among tumour types, only TGCTs and head and neck cancer had the highest number of downregulated piRNAs. Interestingly, low levels of DNA methylation at LINE1 sequences have been reported for TGCTs and head and neck squamous cell carcinoma (HNSCC) represents the second tumour type with the highest LINE1 activation (14, 44).

As a high proportion of piRNAs sequences we identified overlapped with TEs and LINE1 and LINE1-derived antisense transcripts have been primarily identified in the testis (12), we next asked whether the failed silencing and consequent aberrant activity could lead to the expression of oncogenic lncRNAs from LINE1 antisense promoter activity in SEM, a tumour subtype characterised by genome-wide hypomethylation (45). To this end, we analysed piRNAs sequences mapping LINE1-derived lncRNA sequence and identified the pachytene piR-hsa-7221 overlapping the LINE sequence L1PA5 and the antisense transcript CASC9, a lncRNA significantly upregulated in SEM.

The identification of the LINE1-derived CASC9 expression is supported by a recent report in oesophageal cancer demonstrating its regulation through an active LINE1 antisense promoter (46). The functional regulation of CASC9 by piR-hsa-7221 in SEM cells demonstrated that the piRNA binds to the demethylated CASC9-L1PA5 promoter sequence by base paring, independently of the interaction with PIWI proteins. The downregulation of CASC9 was particularly pronounced after transfection of PIWIL1 and PIWIL2, consistent with their association with pachytene piRNAs and with the role of lncRNAs to serve as a source of new piRNAs after their degradation through a positive feedback regulatory loop (18).

It has been shown that only a small fraction (∼1%) of pachytene piRNAs target transposons by sequence complementarity as most pachytene piRNAs are encoded from intergenic regions poor in TE sequences (47). However, blocking of the pachytene piRNA biogenesis leads to reactivation of young and active-transposing TEs, including LINE1 sequences (48). This protective role of pachytene piRNAs is conserved across amniotes, indicating that silencing of TEs is a primary role, which may have evolved to extend their function to post-transcriptional regulation of gene expression (48). In addition, it has been shown that the activity of the mouse Miwi (PIWIL1 in humans) via pachytene piRNA association can target LINE1 transcripts and downregulate the expression of L1ORF1 protein through a slicing activity which leads to double stranded DNA breaks (49). Therefore, this evidence supports or findings demonstrating the regulation of LINE1 sequence by the pachytene piR-hsa-7221, with a direct effect on the expression of the LINE1-derived antisense lncRNA CASC9.

CASC9 has been reported to act as an oncogenic lncRNA in several cancer types, including breast, lung, colorectal, oesophageal, liver, pancreatic ovarian and brain cancers (50).

Its oncogenic activity in these cancer types is demonstrated by enhanced cell proliferation, cell invasion and drug resistance mechanism through activation of proliferative signalling pathways, resistance to apoptosis, induction of EMT, extracellular matrix interaction, and regulation of cell metabolism. These effects are mediated by transcriptional regulation via the recruitment of epigenetic modifiers as well as post-transcriptional regulation via sponging of miRNAs involved in the downregulation of oncogenic transcripts (51). Through loss of function, our data show that CASC9 has a similar oncogenic role in SEM as we measured a reduction of cell proliferation and invasion, as well as improved chemosensitivity after its downregulated expression.

ECM interactions and EMT were among the pathways most significantly affected by CASC9 silencing, which are consistent with the decreased cell migration phenotype. We also found that critical regulators of the cell cycle and developmental signalling pathways were downregulated, which supports the decrease in cell proliferation observed. However, for all these pathways, we did not find any upregulation of known miRNA targets, suggesting these effects are most likely regulated by transcriptional regulation rather than miRNA sponging. Among gene expression signatures, we identified WNT signalling as the most enriched pathway and functional network. This represents a significant finding as WNT signalling plays an important role in germ cell tumours and previous studies have shown its prominent activation and association with tumour relapse and decreased survival rates (52, 53). WNT signalling is also linked to other identified downregulated pathways. For example, WNT can drive EMT and the synthesis of ECM proteins, thus regulating cell invasion (54, 55).

Additional evidence links the WNT pathway to immune evasion in TGCTs (56), as SEM tumours present high infiltration of exhausted immune cells with immunosuppressive features (57). Therefore, downregulation of WNT could also be linked to the downregulation of immune response genes after CASC9 silencing. WNT inhibitors have been evaluated for combination therapy in clinical trials for pancreatic, colorectal and myeloid cancers. These trials have demonstrated safe and effective activity, but no data is available for TGCTs. However, WNT inhibitors have been demonstrated to be very effective in preclinical studies, especially in the treatment of cisplatin-resistant germ cell tumour cell lines (52, 58). Therefore, this study provides new evidence of a therapeutic vulnerability in testicular cancer based on CASC9 expression and paves the way for further research into the development of CASC9 and WNT targeting as a targeted therapeutic strategy for the treatment of SEM.

## Conclusion

Several targeted therapeutics, including protein kinase inhibitors, PARP inhibitors, CDK inhibitors, and immune and epigenetic therapies, have been evaluated in preclinical and clinical studies for testicular cancer. However, none of these drugs have been effective in clinical trials for the treatment of patients who are refractory to cisplatin (59). In this study, we provide new mechanistic insight into SEM carcinogenesis, which is dependent on the expression of the lncRNA CASC9 due to the downregulation of piR-hsa-7221 and associated PIWI proteins. CASC9 plays an oncogenic role by regulating cell proliferation, invasion and chemosensitivity through the reduced expression of genes controlling cell cycle progression, interaction with ECM components and WNT signalling.

Therefore, by providing novel insights into piRNAs expression and their contribution to the epigenetic regulation of LINE1 sequences, CASC9 and its associated gene networks, this study supports the development of new clinical studies for the targeted treatment of SEM.

## Ethics approval and consent to participate

The study was approved by the University of Nottingham CARE committee (Project title: Defining the role on non-coding RNAs in germ cell development and cancer; approval number 158151022). Pair-end RNA-seq data from TGCT patient samples processed were obtained from The Cancer Genome Atlas (TCGA) network after approval (dgGaP Project Approval #12671, Study Accession phs000178).

## Consent for publication

All authors consent to the publication of this article.

## Availability of data and materials

RNA sequencing data are available from GEO (accession number GSE233701; https://0-www-ncbi-nlm-nih-gov.brum.beds.ac.uk/geo/query/acc.cgi?acc=GSE233701). Supporting data are contained in the manuscript as Supplementary Information. Materials are available from the corresponding author upon request.

## Acknowledgements

We acknowledge the kind donation of the TCam-2 cell line by Professor Azim Surani, University of Cambridge. The RNAseq analysis was outsourced to Novogene Co., Ltd (Cambridge, UK).

## CRediT authorship contribution statement

**Ahmad Zyoud:** Investigation, Methodology, Data curation, Formal analysis, Software, Visualization, Writing – original draft, Writing – review & editing. **Ryan Cardenas:** Methodology, Software, Visualization, Validation. **Nabeelah Almalki:** Investigation, Writing – review & editing. **Tinyiko Modikoane**: Investigation, Writing – review & editing. **Mohammed Ageeli Hakami:** Investigation, Validation. **Mansour Alsaleem:** Validation, Writing – review & editing. **Cristina Tufarelli:** Conceptualisation, Writing – review & editing. **Nigel P Mongan:** Data curation, Validation, Formal analysis, Software, Visualization, Supervision, Writing – review & editing. **Cinzia Allegrucci:** Conceptualisation, Investigation, Data curation, Formal analysis, Funding acquisition, Project administration, Resources, Supervision, Writing-original draft, Writing-review and editing.

## Appendix. Supplementary material

**Table S1.** Primers and RNA mimics used in the study.

**Fig. S1.** Overlap of piRNA IDs in piRBase and piRNAdb with testis databases for normal testis (accession numbers ERX2054613, ERX2054614, ERX2054615, GSM2193189) and TGCT datasets. Non-matching piRNAs identified with a different name/annotation are shown in the tables. **A.** Overlap of normal testis samples IDs. **B.** Overlap of TGCTs IDs.

**Fig. S2.** Clustering and correlation of piRNAs expression in normal testis and TGTCs. **A.** Unsupervised hierarchical clustering heatmap of piRNAs in normal testis (blue, n=4), SEM (green, n=58), and NON-SEM (red, n=57). **B.** Spearman correlation-based matrix of piRNA transcripts expression in normal testis (blue, n=4), SEM (green, n=58), and NON-SEM (red, n=57).

**Table S2.** List of piRNAs expressed in different cancer types and overlapping with those expressed in TGCTs.

**Table S3.** List of piRNAs overlapping lncRNA TE and lncRNA sequences.

**Fig. S3. A.** Sequence mapping visualisation showing the overlap of piR-hsa-7221 with the L1PA5 (antisense) and CASC9 (sense) sequences at the chr8:75,324,251-75,324,280 locus (UCSC Genome Browser assembly GRCh38/hg38). **B.** Expression of CASC9 transcript variants in TCam-2 cells. PCR primers were designed to amplify long transcript variants overlapping the piR-hsa-7221 binding sequence and the LINE1-L1PA5 sequence ([1,11,2], [9,11,2]) and [10,11,2], as well as short transcript variants overlapping a LINE1-L1PA4 sequence ([6,6,2] and [6,8,2]).

**Fig. S4.** Log2 Fold change expression of genes regulated by CASC9 silencing. Negative values indicate genes downregulated upon CASC9 silencing, whereas positive values indicate upregulated genes.

**Fig. S5.** Enriched functional protein networks associated with downregulated genes after CASC9 silencing.

## Glossary

FDR: False discovery rate
GEO: Gene Expression Omnibus
miRNA: microRNA
PGC: Primordial germ cell
SEM: Seminoma
NON-SEM: Non-seminomas
TCGA: The Cancer Genome Atlas
TGCT: Testicular germ cell tumour
EMT: epithelial to mesenchymal transition
piRNA: PIWI-interacting RNA
ECM: Extracellular matrix
RNAseq: RNA sequencing
lncRNA: Long non-coding RNA

