## Supplementary material for "Loss of piR-hsa-7221 regulation drives the expression of the LINE1-derived oncogenic lncRNA CASC9 in testicular cancer": Table S1

**Supplementary Table 1. Primers and RNA mimics used in the study.**

| **Gene expression** | **Sequences** |
| --- | --- |
| piR-hsa-7221 | 5’ –TCGCTCACGCTGGGAGCTGTAGACCGGAGC -3’ |
| piR-hsa-7221  stem-loop | 5’-GTCGTATCCAGTGCAGGGTCCGAGGTATTCGCACTGGATACGACGCTCCG -3’ |
| piR-hsa-7221 forward | 5’-GCTCACGCTGGGAGCTGTAGAC-3’ |
| piR-hsa-7221 reverse | 5’ -CCAGTGCAGGGTCCGAGGTA-3’ |
| PIWIL1 forward | 5’-AAGCCAGAAGACTCCGTTCA-3’ |
| PIWIL1 reverse | 5’-TCACATCCTCTCCATTCCGG-3’ |
| PIWIL2 forward | 5’-CGTGGCAAGCATCAATCTCA-3’ |
| PIWIL2 reverse | 5’-TTGGCCATCAGACACTCCAT-3’ |
| PIWIL4 forward | 5’-GCTTGTCCTGTTTCACCCAG-3’ |
| PIWIL4 reverse | 5’-TAGGTGATCTCGGTGCCATC-3’ |
| CASC9-all variants forward | 5’-TGTGCTAGCAATCAGCGAGA-3’ |
| CASC9-all variants reverse | 5’-GCCAGGTGTTGTTCTGCTATC-3’ |
| CASC-V1 forward  Variant [1,11,2]  Variant [9,11,2]  Variant [10,11,2] | 5’-CGTCTTCTGCGTCGCTCAC-3’ |
| CASC9-V2 forward  Variant [6,6,2]  Variant [6,8,2] | 5’- CCCAGTTGGAGCTTCCTG-3’ |
| CASC9-V1/2 reverse Variants | 5’-AGTGCCAATGACTCTCCAGC-3’ |
| U6 snRNA forward | 5’-CTCGCTTCGGCAGCACA-3’ |
| U6 snRNA reverse | 5’-AACGCTTCACGAATTTGCGT-3’ |
| ACTB forward | 5’-ATAGCAACGTACATGGCTGG-3’ |
| ACTB reverse | 5’-CACCTTCTACAATGAGCTGC-3’ |
| **Plasmids** | **Sequences** |
| LINE1-psiCHECK2 forward | 5’- TTCGCTCGAGACCGGAGGAGCCAAGATGGC-3’ |
| LINE1-psiCHECK2 reverse | 5’- TTTGCGGCCGCCGAAAAGCGCAATATTCGGGT-3’ |
| PIWIL1 forward | 5’-TTTACTAGTACCATGACTGGGAGAGCCCGA-3’ |
| PIWIL1 reverse | 5’-AGCGGAATTCTTAGAGGTAGTAAAGGCGGTT-3’ |
| PIWIL2 forward | 5’- TTTACTAGTACCATGGATCCTTTCCGACCA -3’ |
| PIWIL2 reverse | 5’- ATCCGAATTCTCACAGGAAGAACAGGTTCT -3’ |
| PIWIL4 forward | 5’- TTTACTAGTACCATGAGTGGAAGAGCCCGAG-3’ |
| PIWIL4 reverse | 5’- AGTTAGCGCTTCACAGGTAGAAGAGATGGTT-3’ |
| **DNA methylation** | **Sequences** |
| L1Methyl-proximal forward | 5’-GAGATTATATTTTATATTTGGTTTAGAGGG-3’ |
| L1Methyl-proximal reverse | 5’- AACTATAATAAACTCCACCCAATTC-3’ |
| L1Methyl-distal forward | 5’- TTATTAGGGAGTGTTAGATAGTGGG-3’ |
| L1Methyl-distal reverse | 5’- TTGTTTAGGTTTGTTTAGGTA-3’ |
| **Mimics and siRNA** | **Sequences and assays** |
| piR-hsa-7221 | 5’ – UCGCUCACGCUGGGAGCUGUAGACCGGAG [mC]-3’ (Sigma-Aldrich) |
| RNA scrambled | 5’ – UUGGUGCUCUUCAUCUUGUUG -3’ (Sigma-Aldrich; MISSION®, SHC002) |
| CASC9 siRNA | Silencer® Select assay ID: n544070 |
| CASC9 siRNA | Silencer® Select assay ID: n544071 |
