## Supplementary figures and images for "Loss of piR-hsa-7221 regulation drives the expression of the LINE1-derived oncogenic lncRNA CASC9 in testicular cancer"

### Supplementary Figures

**Fig. S1**

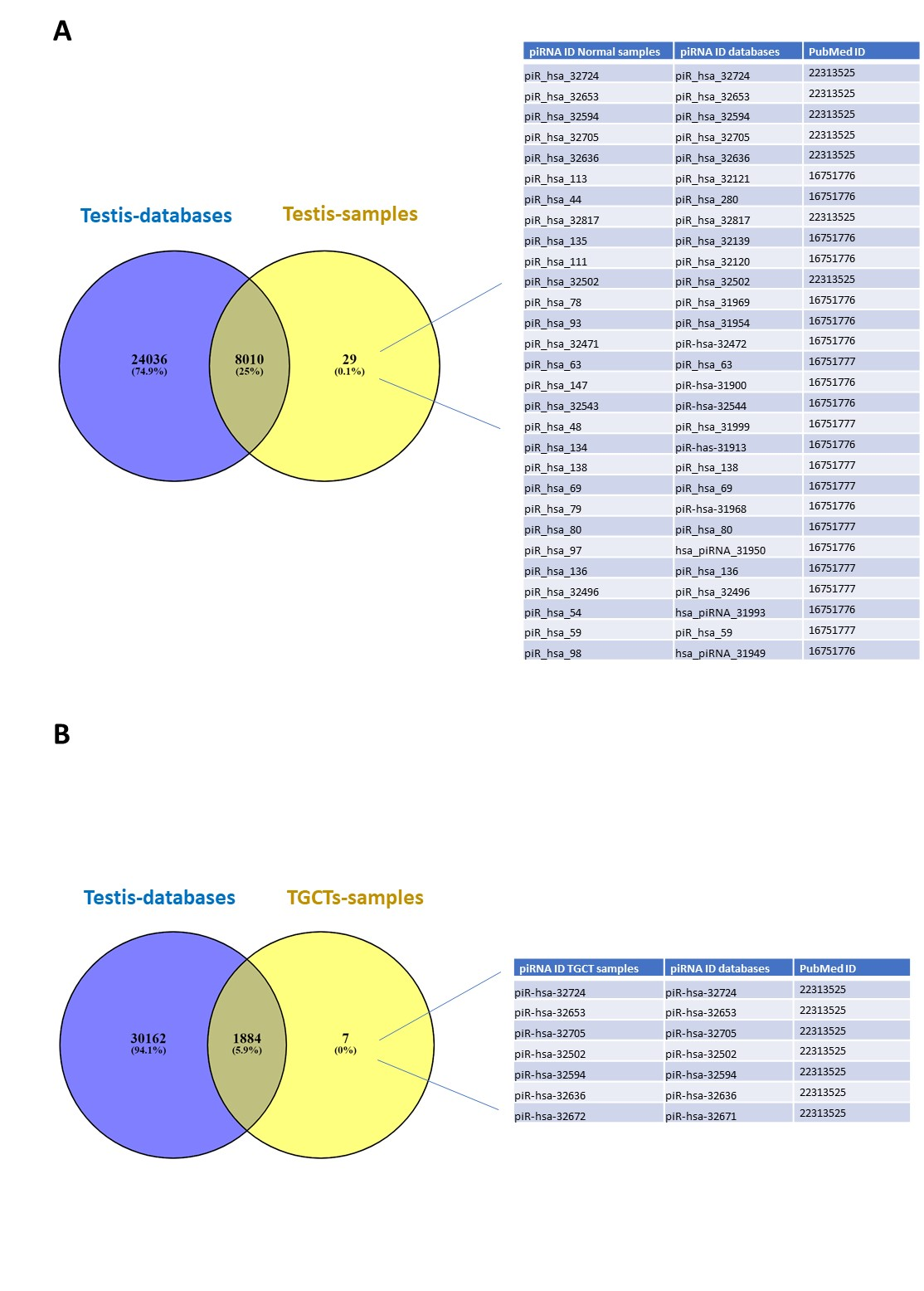

**Fig. S2**

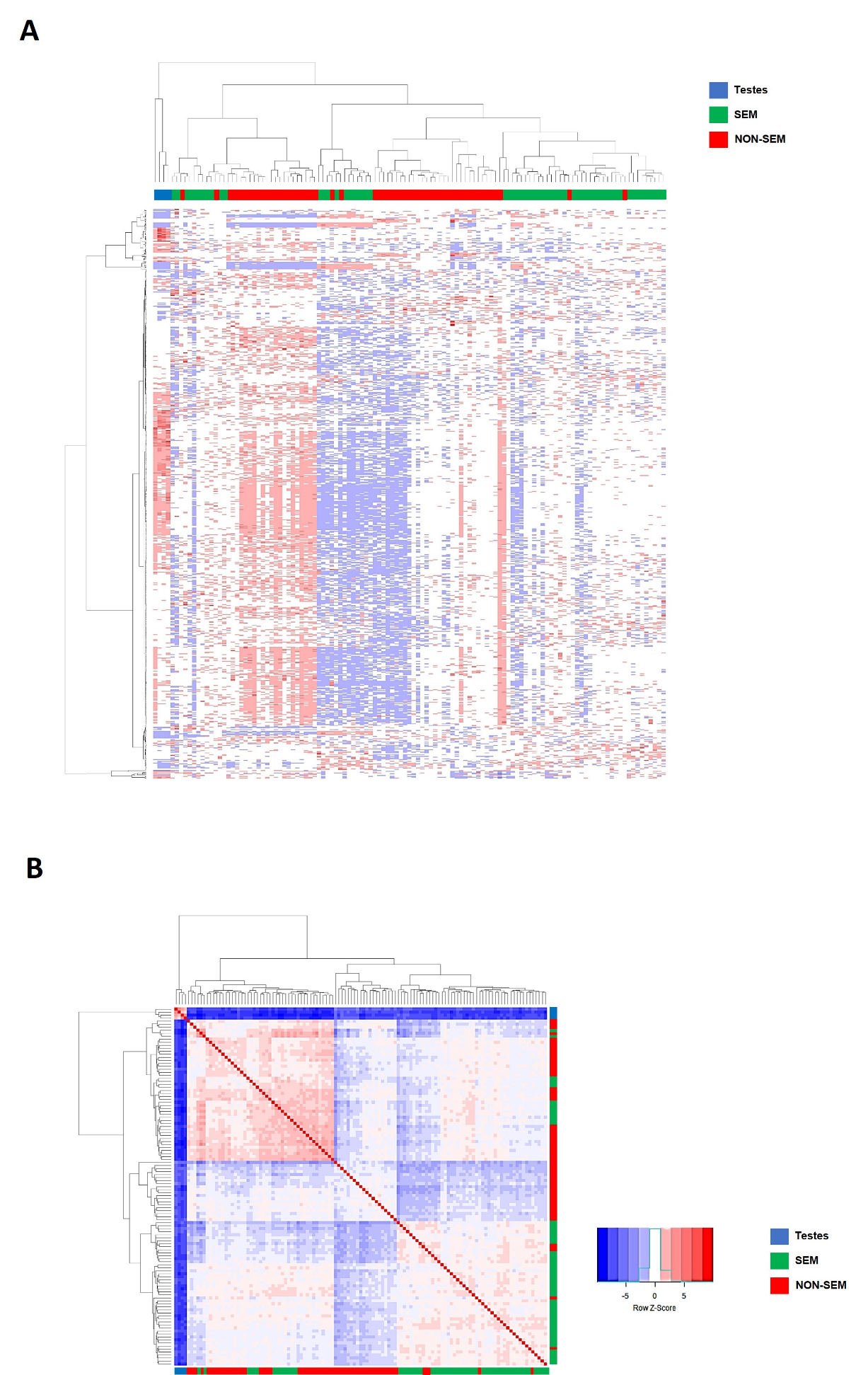

**Fig. S3**

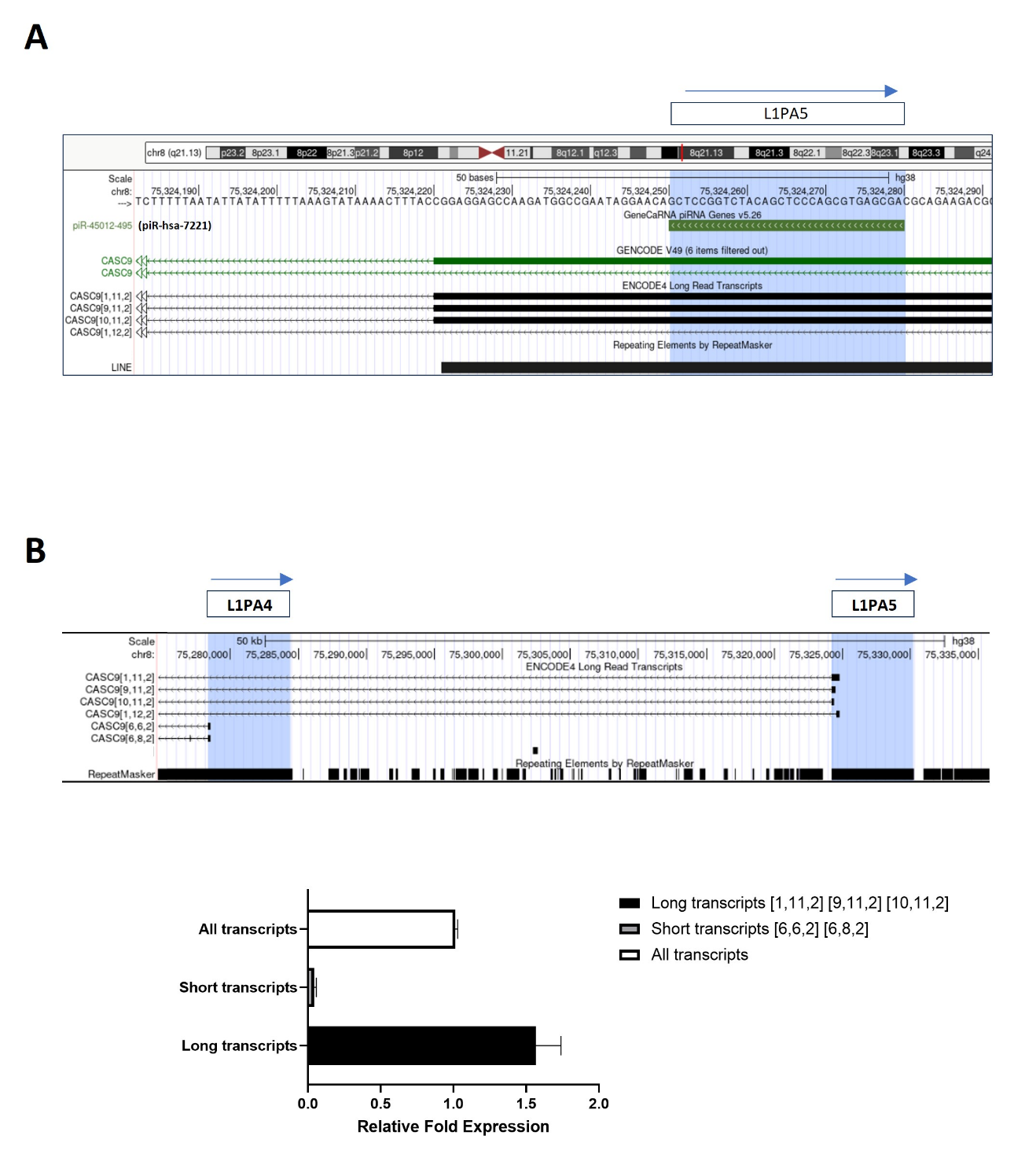

**Fig. S4**

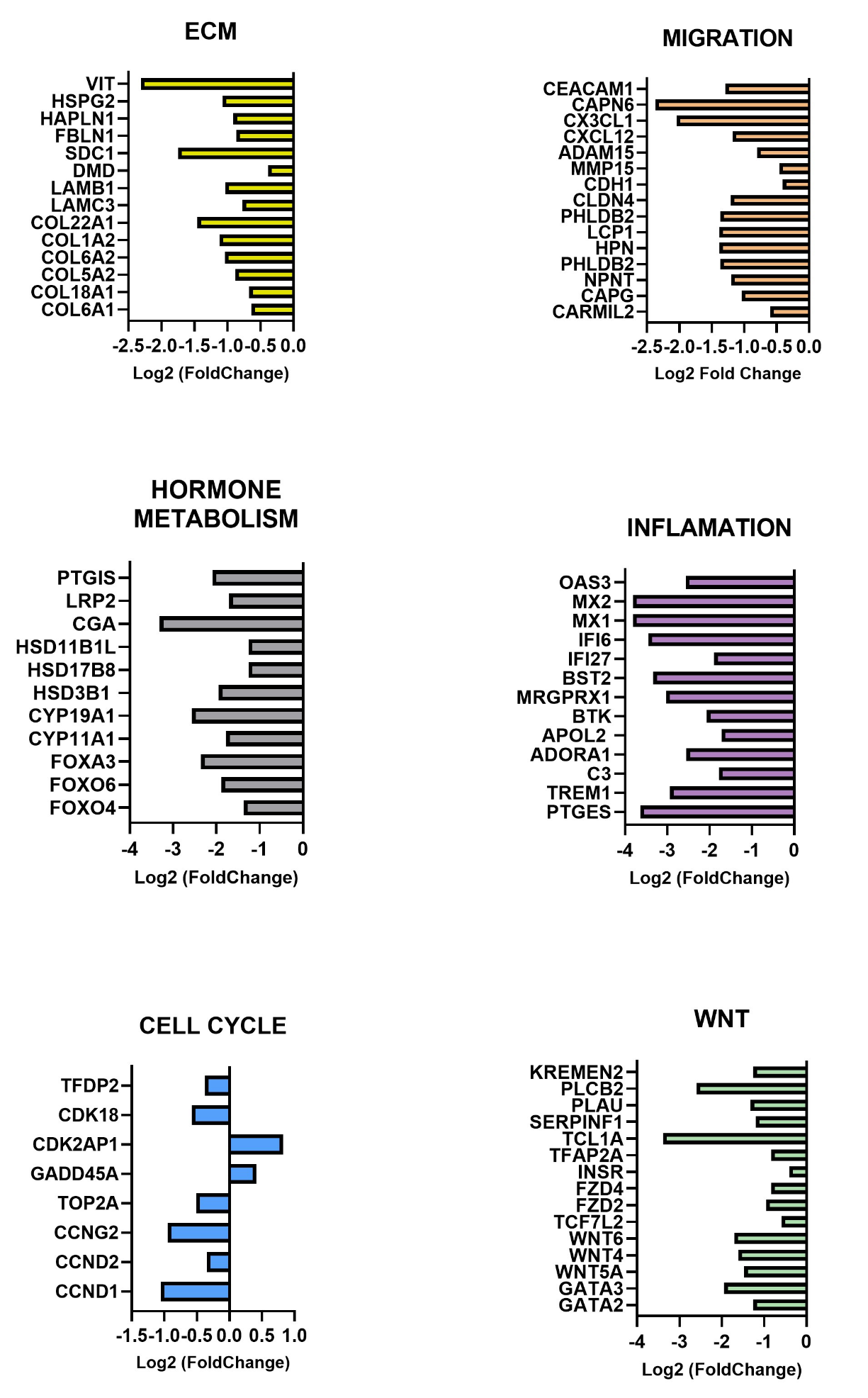

**Fig. S5**

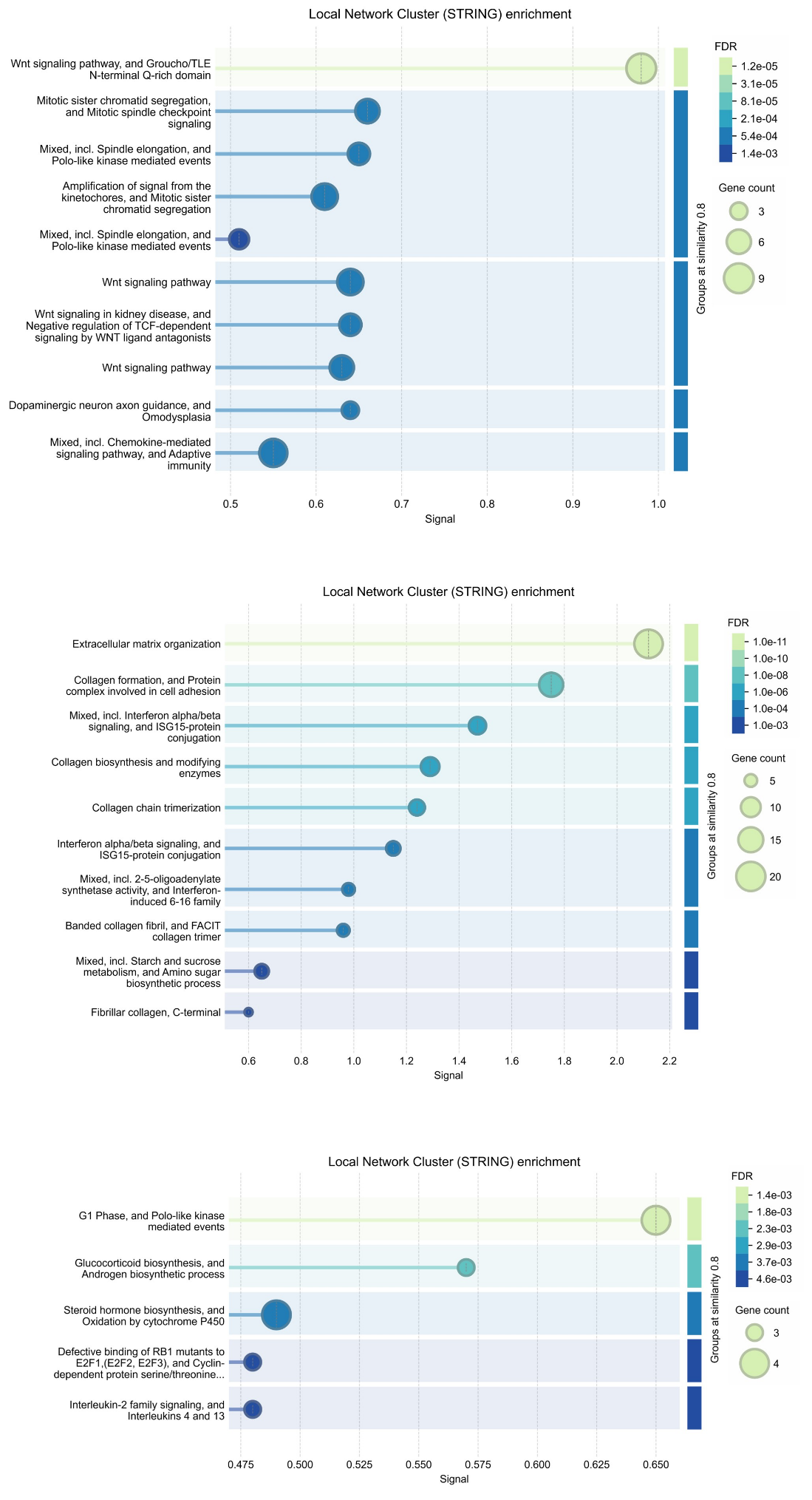
